# Meiotic drive does not impede success in sperm competition in the stalk-eyed fly, *Teleopsis dalmanni*

**DOI:** 10.1101/2022.09.09.507334

**Authors:** Sadé Bates, Lara Meade, Andrew Pomiankowski

## Abstract

In male X-linked meiotic drive systems, the driver causes degeneration of Y-bearing sperm, leading to female-biased offspring sex ratios. This potentially leads to a two-fold transmission advantage to drive chromosomes. However, drive-bearing sperm often do poorly in sperm competition, limiting their ability to spread. We use the stalk-eyed fly, *Teleopsis dalmanni*, to investigate the success of the X-linked *Sex Ratio* (SR) meiotic drive system. In this species, polyandrous matings, where a female mates with multiple males, are common. Recent findings demonstrate SR males transfer the same numbers of viable sperm as wildtype (ST) males during mating, implying that they do not necessarily have reduced fertility under sperm competition. Reciprocal mating trials were performed to measure the success of SR and ST sperm in double mated females, with either a SR or ST male mated first followed by a male of the alternative genotype. There was no significant difference in the number of offspring sired by SR and ST males. This equivalence held regardless of whether the SR male mated first or second. We show these results are consistent with previous studies that suggested SR male sperm do poorly in sperm competition. Future experiments will determine whether the competitive ability of SR males is maintained under higher stress conditions likely to be experienced in nature, in which females mate repeatedly with multiple males. The results from the current study helps to explain the high meiotic drive frequency of around 20% in wild populations in this species.

**Impact Summary:** Meiotic drive genes are selfish genetic elements that distort Mendelian patterns of inheritance to bias transmission in their favour. We use the stalk-eyed fly, *Teleopsis dalmanni*, to investigate the fitness effects associated with a meiotic drive gene called *Sex Ratio* (SR), which is linked to the X chromosome. In males, SR destroys Y-bearing sperm, meaning only X-bearing sperm are viable, and females who mate with drive males sire all-female broods. This confers a two-fold transmission advantage to the SR gene, as it is transmitted to all offspring.

We recently discovered that drive males have evolved compensatory mechanisms to cope with the sperm destruction caused by meiotic drive. They have greatly enlarged testes, allowing them to produce more sperm. When drive males mate with females, they deliver as many sperm and sire as many offspring as wildtype males. Building on this finding, we measured how drive male sperm performs against sperm from a non-carrier male in sperm competition – where the sperm from different males compete to fertilise an egg. Double mating trials were performed, where a single female was mated once to a drive and once to a non-carrier male. By genotyping offspring, we show that the number of offspring sired by the drive male was not different from the number sired by the non-carrier competitor.

These findings contrast with those in other species. Typically, drive males do poorly in sperm competition and their spread is severely restricted by sperm competition. In stalk-eyed flies, female multiple mating with many males is the norm, but this does not appear to inhibit the fertility of drive males. The success of drive under sperm competition helps to explain the high frequency of drive around 20% in natural populations of *T. dalmanni*.

## Introduction

Meiotic drive causes the unequal transmission of genes in the next generation (Gershenson, 1928; Sandler & Novitski, 1957). In the extreme, the driver entirely excludes wildtype alleles and is transmitted to all offspring, violating Mendelian laws of segregation (Searle & de Villena, 2022; Wolf, Ferguson-Smith, & Lorenz, 2022). This is most potently seen in X-linked drivers, which are common among Diptera species and lead to dysfunction of Y-bearing sperm and the production of female-only broods (Policansky, 1974; Newton, Wood, & Southern, 1976; James & Jaenike, 1990; Hurst & Pomiankowski, 1991; Presgraves, Severance, & Wilkinson, 1997; Jiggins, Hurst, & Majerus, 1999). Such a significant transmission advantage could potentially lead to population extinction due to the lack of males (Hamilton, 1967; Hatcher, Taneyhill, Dunn, & Tofts, 1999; Mackintosh, Pomiankowski, & Scott, 2021). However, the fitness costs associated with carrying drive genes often result in negative frequency-dependent selection that limits their spread (Lindholm *et al*., 2016; Finnegan *et al*., 2019). As drive genes are typically located in low frequency inversions subject to weak selection, deleterious effects likely accumulate after their origin, contributing to restrictions to their population frequency (Dyer, Charlesworth, & Jaenike, 2007; Reinhardt *et al*., 2014; Navarro-Dominguez *et al*., 2022). In turn, these fitness costs favour the evolution of modifiers that suppress transmission distortion, which occurs both on the wildtype and drive chromosome as well as elsewhere in the genome (Keais, Lu, & Perlman, 2020).

One factor that has a strong impact on the spread of meiotic drive genes is reduced fertility (Zanders & Unckless, 2019). In males with complete drive, all wildtype gametes are lost and the number of mature sperm is reduced. For example, in the well-studied SD system in *D. melanogaster*, males heterozygous for drive have reduced fertility in non-competitive matings due to lower sperm production (Hartl, 1969). In addition, pleiotropic “collateral damage” caused by drive may reduce the activity or number of mature drive sperm, leading to poor outcomes when sperm from drive males compete for fertilizations with sperm from wildtype males (Price & Wedell, 2008). This deficit is likely to be prominent in insects that possess reproductive organs specialised for long-term storage of viable sperm, increasing interactions between ejaculates and thus sperm competition (Parker, 1970). Evidence from sperm competition studies of sex-linked meiotic drive systems in *Drosophila* species supports this prediction. In *D. pseudoobscura*, SR drive males sire fewer offspring than standard males in double mating trials (Price *et al*., 2008). Drive males have a disproportionally lower success as the first (P1) and second male (P2) against wildtype males, reflecting a lack of sperm competitiveness both in their ability to resist other sperm or to displace sperm already in storage (Price *et al*., 2008). A similar pattern occurs in *D. simulans* with reduced success in P1 and P2 for drive males, and preferential drive male sperm ejection from the female reproductive tract even without competition from the second male’s sperm (Atlan, Joly, Capillon, & Montchamp-Moreau, 2004; Angelard, Montchamp-Moreau, & Joly, 2008). It has been suggested that increased female polyandry evolves to undermine the success of drive sperm and an experimental evolution study in *D. pseudoobscura* and a double mating experiment in *D. recens* support this possibility, linking the frequency of drive with the rate of multiple mating (Haig & Bergstrom, 1995; Zeh & Zeh, 1997; Price, Hodgson, Lewis, Hurst, & Wedell, 2008; Courret, Chang, Wei, Montchamp-Moreau, & Larracuente, 2019; Dyer & Hall, 2019).

In this paper we show that these fertility deficits do not generalise to the X-linked SR meiotic drive in the stalk-eyed fly *Teleopsis dalmanni*. Stalk-eyed fly females store sperm in the spermathecae (long-term storage organs) after mating, before sperm migrate to the ventral receptacle, where they are individually packaged into pouches prior to release into the oviduct for fertilisation of mature eggs. In several stalk-eyed fly species, the main mode of sperm competition is sperm mixing, rather than male precedence (Lorch, Wilkinson, & Reillo, 1993; Wilkinson, Kahler, & Baker, 1998; Corley *et al*., 2006; Bellamy, 2012). Under this model, the share of each male’s paternity is dependent on the number and quality of sperm he transfers per ejaculate. It was predicted that drive males should be poor competitors due to the degeneration of Y-bearing sperm and the deleterious effects associated with carrying a driver gene. Double mating trials appear to confirm this as drive (SR) males sired fewer offspring than wildtype (ST) males (Wilkinson & Fry, 2001; Wilkinson, Johns, Kelleher, Muscedere, & Lorsong, 2006). However, several factors raise concerns about a simplistic interpretation of these findings. The Wilkinson and Fry (2001) study performed an inter-population cross of Malaysian and Thai flies, and outcomes could have been confounded by genetic incompatibility, which has been predicted to be stronger for drive chromosomes (Hurst & Pomiankowski, 1991). In addition, in the absence of genetic markers offspring were distinguished by eye using heritable variation in leg colour, for which an error-rate is unknown. Neither of these matters affect the experimental design of a second study which again reported lower SR male paternity (Wilkinson, Johns, Kelleher, Muscedere, & Lorsong, 2006). However, this effect was limited to single-parent broods, which were less frequently fathered by SR males. There was no difference in SR and ST paternity in mixed paternity broods. In addition, this experiment only considered the competitive ability of SR males when mating second. This means that defensive traits of SR sperm and ejaculate were not assessed, so it is unclear whether the lack of success of SR males is general or limited to lower sperm precedence when mating second.

Whilst these points prompt concern over the robustness of the previous results, a more profound challenge arises from recent findings that SR males transfer similar numbers of sperm per ejaculate (Meade *et al*., 2019; Meade, Finnegan, Kad, Fowler, & Pomiankowski, 2020). This was measured in females both in the spermathecae as well as the movement of sperm to the ventral receptacle after matings with SR or ST males, as well as after up to three sequential matings by a single male (Meade *et al*., 2019). Furthermore, when egg counts were used to measure fertility after single matings, it did not differ for females mated to SR or ST males (Meade, Finnegan, Kad, Fowler, & Pomiankowski, 2020). Dissection of adult SR males reveals that they have greatly enlarged testes which allow sperm delivery and fertility to be maintained despite the destruction of sperm caused by meiotic drive (Meade *et al*., 2019; Meade, Finnegan, Kad, Fowler, & Pomiankowski, 2020). This challenges the conventional view that SR males are weak competitors, and specifically, the finding of a competitive deficit of SR males in double mating trials.

Here, we studied whether SR males are at a disadvantage during sperm competition. The paternity of SR males was measured by performing reciprocal double mating trials in which the SR male mated first followed by the ST male, or vice versa. This allowed an assessment of the SR male’s success in both the offensive and defensive capabilities. The offspring arising from these trials were collected and genotyped at the larval stage to determine the proportion of offspring sired by SR males. This enabled the study to avoid confounds in paternity share relating to egg to adult viability differences, which have recently been shown to disadvantage SR-carrying larvae (Finnegan *et al*., 2019).

## Methods

### Stock populations

Flies for the standard stock (ST-stock) population carry only the wildtype X chromosome (X^ST^). They were collected (by S. Cotton and A. Pomiankowski) in 2005 from the Ulu Gombak valley, Peninsular Malaysia (3°19′N 101°45′E). They have since been maintained in high density cages (> 200 individuals) to minimise inbreeding and are regularly monitored to ensure they do not contain meiotic drive.

The meiotic drive stock (SR-stock) population is composed of females that are homozygous for a sex-ratio distorting X chromosome (X^SR^). They were derived from flies collected in 2012 (by A. Cotton and S. Cotton) from the same location as the ST-stock. X^SR^/Y males produce 100% female offspring due to transmission distortion. The X^SR^ female stock is maintained by crossing X^SR^/X^SR^ females with X^ST^/Y males to produce X^SR^/Y drive males, who are then mated to the X^SR^/X^SR^ females to generate the next generation of the SR-stock females. The outcrossing to ST males from the ST-stock ensures that the two stocks only differ in their X chromosomes and are homogenised for autosomal content.

Both stock populations were kept at 25 °C, with a 12:12 h dark:light cycle and fed puréed sweetcorn twice weekly. Fifteen-minute artificial dawn and dusk periods were created by illumination from a single 60W bulb at the start and end of the light phase.

### Experimental populations

Experimental ST (X^ST^/Y) and SR males (X^ST^/Y) were drawn from the ST-stock and SR-stock, respectively. They were housed separately in cages of ∼50 individuals until sexually mature, in groups of similar age (6-8 weeks). ST-stock females were added to these cages at an equal sex ratio for > 3 days to allow males to mate. The females were then removed and discarded. Experimental males were then kept in single-sex groups for a further 3-6 days to allow their accessory glands to return to full size (Rogers, Chapman, Fowler, & Pomiankowski, 2005).

Experimental ST females (X^ST^/X^ST^) were drawn from the ST-stock. All experimental females were virgins, 6-8 weeks old, and had reached sexual maturity (Baker *et al*., 2003). ST females were anaesthetised on ice and eyespan was measured (see below), in order to exclude small flies and limit variation in size and fecundity that could influence sperm allocation strategies in males (Cotton, Cotton, Small, & Pomiankowski, 2015). Only large females with eyespan > 5.4 mm were used in mating trials (range 5.4 – 5.8 mm).

### Sperm competitiveness of SR and ST males

Mating trials were conducted to measure the competitiveness of SR and ST males. On the day preceding each assay, experimental females were housed singly in 500ml clear plastic containers with a moist cotton wool base. On the trial day, a single male was added to each container approximately 15 minutes after dawn, as this is the period during which mating is most likely (Chapman, Pomiankowski, & Fowler, 2005). Males were allowed to mate, defined as copulation lasting >30s, which is sufficient for sperm transfer (Cotton, Cotton, Small, & Pomiankowski, 2015). The mating duration was recorded. If no mating was observed after 15min, the male was moved to a new container with a new female. If mating still did not occur after a further 15min, the male was discarded. The original unmated female was used again and placed with another male. If this did not result in a copulation after 15min, the female was discarded.

A second mating was performed 24hrs later, following the same protocol. If the female failed to mate on the second day she was discarded. Females were mated either to a SR male followed by a ST male, or a ST male followed by a SR male. Once females had been double mated, the containers were lined with a fresh moist cotton wool base and 1tsp puréed sweetcorn, which was collected and renewed every 2-3 days for 2 weeks. This kept larval density low, maximising survival. Bases were stored in Petri dishes at 25ºC. In total, 72 females were successfully mated twice: 36 to a SR male first and 36 to an ST male first. For ease, these matings were carried out in two batches, one week apart.

After mating, experimental males were removed and frozen and their eyespan and thorax measured under a Leica microscope using ImageJ (v1.46; Schneider *et al*., 2012). Eyespan was defined as the distance between the outer tips of the eyes (Hingle, Fowler, & Pomiankowski, 2001). Thorax length was defined as the distance ventrally from the anterior tip of the prothorax along the midline to the joint between the metathoracic legs and the thorax (Rogers, Denniff, Chapman, Fowler, & Pomiankowski, 2008).

### Progeny genotyping

Petri dishes were examined for larvae one week after collection and then daily. Once larvae had developed to be large enough to seen by eye, they were transferred to a 96-well plate. Each well of the plate contained 100µL digestion solution (20mM EDTA, 120mM NaCl, 50mM Tris-HCL, 1% SDS, pH 8.0) and 4µL proteinase K (10mg ml−1). A standard protocol was adapted to extract and purify DNA from larvae (see SI for details; protocol from Burke et al., 1998). The X-linked INDEL marker *comp162710* was used to identify offspring of ST and SR fathers, due to its reported accuracy in determining phenotype (>90%; Meade *et al*. 2019). X^ST^ carries a large allele (286 bp), whereas X^SR^ carries a small allele (201 bp).

### Statistical methods

To test if mating order or genotype affected the number of offspring sired by each male, P1:P2 offspring (the number of offspring sired by P1 relative to the number of offspring sired by the P2 male) or ST:SR offspring (the number of offspring sired by the ST relative to the number of offspring sired by the SR male) were fitted as the response variable in Generalised Linear Models (GLMs) with a binomial error distribution. 15 females that had produced fewer than 10 genotyped offspring were excluded from the analysis. As the GLMs were over-dispersed, a quasi-binomial error distribution was used. The number of larvae collected and the batch were assessed as potential confounding variables.

The effect of male thorax length (a proxy for body size) and relative eyespan (the variation in eyespan after controlling for thorax length) were considered in the analysis. Both traits are strongly condition-dependent and indicators of male genetic and phenotypic quality (David, Bjorksten, Fowler, & Pomiankowski, 2000; Cotton, Cotton, Small, & Pomiankowski, 2015; Howie, Dawson, Pomiankowski, & Fowler, 2019). Whether these male trait sizes differed between genotypes was tested by fitting thorax length and relative eyespan as the response variable in linear models. In addition, differences in mating duration by mating order and genotype were tested as a response variable in linear models, along as confounding variables sas fixed effect in GLMs on the number of offspring sired by each male. Full statistical analyses are reported in the Supplementary Material.

## Results

### Male fertility

In total, 62 females were reciprocally mated to males of each genotype and 47 females had ≥10 of their offspring successfully genotyped to determine the distribution of paternity (23 P1 SR—P2 ST and 24 P1 ST—P2 SR).

The proportion of offspring sired by the P2 male was 0.547 ± 0.046 (mean ± se) and there was no effect of mating order on the number of offspring sired by each male (F_1,43_ = 0.987, *P =* 0.326; Figure 1). The proportion of offspring sired by the SR male was 0.543 ± 0.046 and there was no effect of genotype on the number of offspring sired by each male (F_1,43_ = 0.069, *P =* 0.793; Figure 2). Nor was there an effect of genotype when the data was split in two halves, either with the SR male in the P1 role (F_1,44_ = 0.152, *P* = 0.699) or in the P2 role (F_1,44_ = 0.016, *P* = 0.899. The main effects of order and genotype were unchanged when the number of larvae collected or batch number were included as covariates (*P* > 0.05; see **SI**).

**Figure 1:**
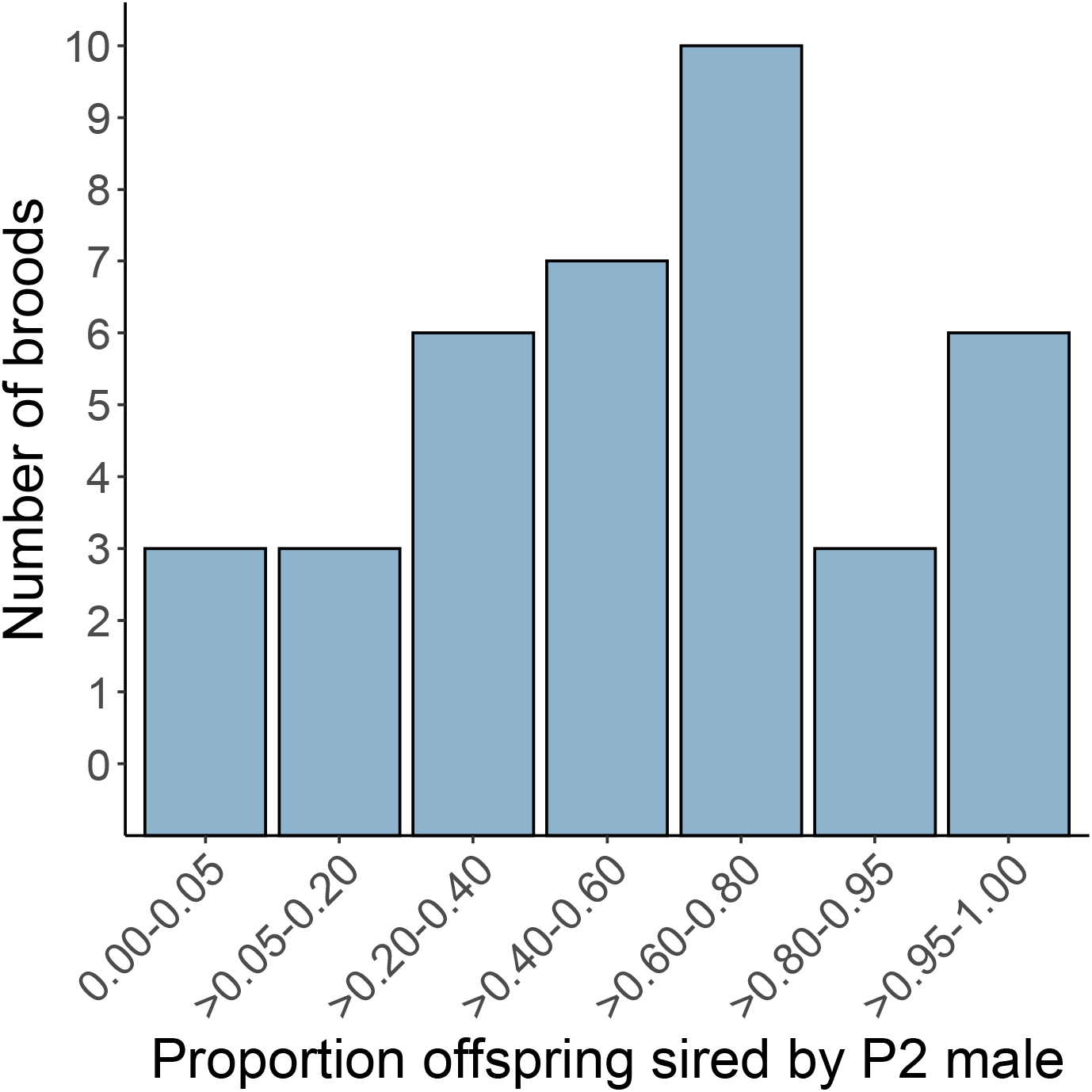
The distribution of the proportion of offspring sired by the second male, P2, is shown per brood.

**Figure 2:**
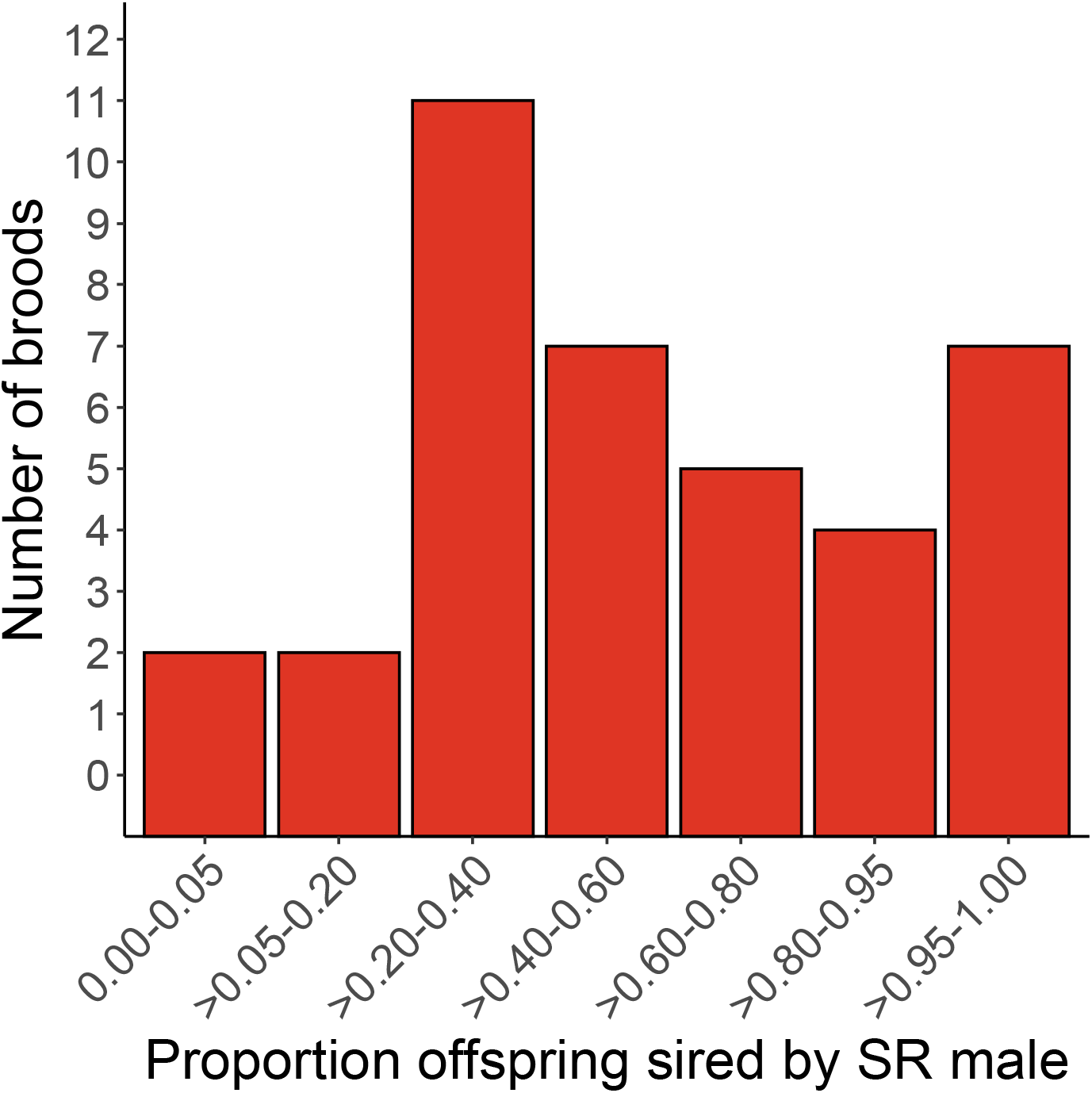
The distribution of the proportion of offspring sired by the SR male is shown per brood.

In 11 of the 47 cases, one male sired more than 0.95 of the offspring, with no difference between male mating position (4 sired by the P1 male, and 7 sired by the P2 male, F_1,9_ = 0.986, *P =* 0.351) or male genotype (8 sired by the SR male and 3 were sired by the ST male, F_1,9_ = 0.841, *P =* 0.383). When these extreme cases were excluded, there was still no effect of mating order (F_1,32_ = 0.094, *P =* 0.761) or male genotype (F_1,32_ = 0.589, *P =* 0.448) on the number of offspring sired.

### Male trait size and mating duration

Thorax length was smaller in SR than ST males (mean ± se, SR = 2.194 ± 0.025 mm, N = 46, ST = 2.285 ± 0.026 mm, N = 46; F_1,88_ = 5.644, *P =* 0.02). Eyespan is strongly colinear with thorax (F_1,88_ = 151.0, *P* < 0.001) and was likewise smaller in SR males (SR = 7.329 ± 0.116 mm, ST = 7.844 ± 0.122 mm; F_1,88_ = 8.868, *P =* 0.004; Figure S1). However, relative eyespan did not differ between genotypes (F_1,87_ = 3.002, *P =* 0.087; Figure S2). As thorax length differed between genotypes, it was added as a covariate, but there was still no effect of mating order (F_1,39_ = 0.748, *P* = 0.392) or male genotype (F_1,39_ = 0.168, *P* = 0.684) on the number of offspring sired by each male.

Mating duration did not differ between mating positions (mean ± se, P1 = 64 ± 3 sec, P2 = 76 ± 3 sec; F_1,90_ = 1.38, *P* = 0.243) or genotypes (ST = 61 ± 3 sec, SR = 79 ± 10 sec, F_1,90_ = 2.66, *P* = 0.106). Mating duration did not affect the number of offspring sired by the P1 (F_1,42_ = 3.474, *P* = 0.069) or P2 males (F_1,42_ = 0.048, *P* = 0.827). Nor did mating duration did not affect the number of offspring sired by the SR males (F_1,42_ = 0.188, *P* = 0.667) but ST males with a shorter mating duration sired a greater number of offspring (F_1,42_ = 4.292, *P* = 0.044). Given this finding, the mating durations of the two males were added as covariates, but there was still no effect of mating order (F_1,41_ = 0.756, *P* = 0.390) or male genotype (F_1,41_ = 0.001, *P* = 0.980) on the number of offspring sired.

## Discussion

Our study provides no support for the idea that males carrying X-linked meiotic drive are at a disadvantage under sperm competition due to sperm loss and other deleterious effects of meiotic drive on sperm function (Courret, Chang, Wei, Montchamp-Moreau, & Larracuente, 2019; Verspoor, Price, & Wedell, 2020). Here, the paternity of SR males did not differ from ST males overall, nor in the P1 or P2 positions considered separately. This challenges the general pattern which has been reported across the Diptera (Policansky, 1974; Newton, Wood, & Southern, 1976; James & Jaenike, 1990; Hurst & Pomiankowski, 1991; Presgraves, Severance, & Wilkinson, 1997; Jiggins, Hurst, & Majerus, 1999; Price *et al*., 2008; Dyer & Hall, 2019). It is also in opposition to previous evidence of lower drive male paternity in stalk-eyed fly double-mating experiments, the limitations to which were discussed in the Introduction (Wilkinson and Fry, 2001; Wilkinson *et al*. 2006). Our results are robust to a number of potential confounding factors: matings were performed between flies from the same population, offspring paternity was assessed using highly accurate genetic markers, larvae were used to assess paternity which reduces the impact of lower egg-adult viability in SR females and double matings were carried out with SR males in the first and second mating position to reliably assess sperm precedence. Furthermore, the findings here align with those of Meade *et al*. (2019, 2020), who showed that sperm numbers transferred to females and the resulting fertility do not differ in single matings by SR and ST males.

Our results do not invalidate previous findings, which likely reflect genuine experimental differences. The study of Wilkinson and Fry (2001) was carried out on the closely related species *T. whitei*, which also carries X-linked SR meiotic drive that is thought to have evolved prior to the divergence of these two species (Presgraves, Severance, & Wilkinson, 1997; Panhuis & Wilkinson, 1999). Genetic markers for drive have not been identified in *T. whitei* (G. S. Wilkinson, personal communication), implying a small inversion is associated with drive in this species, unlike the multiple inversions that cover most of the *T. dalmanni* SR X chromosome (Wilkinson, Johns, Kelleher, Muscedere, & Lorsong, 2006; Christianson, Brand, & Wilkinson, 2011; Reinhardt *et al*., 2014; Paczolt, Reinhardt, & Wilkinson, 2017). This means that few X-linked genes are in linkage disequilibrium with those that control drive, limiting the possibility of compensatory testes enlargement in *T. whitei*, leaving drive males with lowered sperm production and reduced fertility under sperm competition. The second study of Wilkinson *et al*. (2006) used a similar double mating design in *T. dalmanni* (although only with SR males in the P2 role). As in this study, it reported no difference between SR and ST success in mixed paternity broods. However, there was an excess of ST male success in single-parent broods, where only one male fathered offspring. There were 14 single-parent broods of 41 typed, with only 3 derived from the SR male. Whilst this rate is similar to that reported here (11 single-parent broods of 47 typed), there was no association with genotype, with a non-significant excess of SR males (9 derived from the SR male). Combining across these two studies, we conclude that there can be little confidence that there is a real deficit in SR male single-parent broods, say for example because of a higher failure to fully form a spermatophore.

In line with earlier work on sperm competition in stalk-eyed flies, there was no effect of mating order on paternity, suggesting that the sperm of the first and second male simply mix and there is no sperm precedence in *T. dalmanni* (Wilkinson & Fry, 2001; Corley *et al*., 2006; Bellamy, 2012). Corley *et al*. (2006) found evidence of a trimodal P2 distribution, centred around equal paternity as well as a strong bias to either the first or second male (double mating with ST males). This contrasts with the multimodal distribution shown here (Figure 1). The difference could be due to the multiple mating design used by Corley *et al*. (2006), in which each female was mated three times with the first and second males. A trimodal pattern was also reported in a double mating design in the distantly related South African stalk-eyed fly species *Diasemopsis meigenii*, where extreme paternity bias was explained by the failure of sperm transfer after a copulation (Bellamy, 2012). Whatever the explanation, none of these studies support the idea of a competitive advantage associated with mating position in stalk-eyed flies.

In this study of *T. dalmanni*, sperm competition was assessed under low-stress conditions. Virgin females were mated to two males separated by a 24 hour period. Experimental males were not virgins, but had been kept for several days in single-sex groups. The objective was to assess SR and ST males under standardised conditions as a first step to understand how SR males perform under sperm competition. In the wild, competitive conditions are more complex. Males form leks with multiple females at dusk, and then mate in a short period at dawn before dispersal, with occasional matings interspersed during day light hours (Wilkinson, Presgraves, & Crymes, 1998; Chapman, Pomiankowski, & Fowler, 2005; Cotton, Small, Hashim, & Pomiankowski, 2010; Cotton, Cotton, Small, & Pomiankowski, 2015). Females mate repeatedly in a life span that can extend over several months (Wilkinson, Presgraves, & Crymes, 1998; Reguera, Pomiankowski, Fowler, & Chapman, 2004). Multiple matings are required as males transfer low numbers of sperm per ejaculate (Wilkinson, Amitin, & Johns, 2005; Rogers, Grant, Chapman, Pomiankowski, & Fowler, 2006; Meade *et al*., 2019), several matings are needed to reach maximum fertility (Baker *et al*., 2001) and sperm usage leads to a quick drop in female fertility over time (Wilkinson, Kahler, & Baker, 1998; Meade *et al*., 2017). Future experiments need to assess the success of single SR and ST male matings in females with a background of multiple mating, closer to the conditions found in nature. There may be differences when female sperm storage organs are saturated compared to the situation with double mating when females are below maximal fertility (Baker *et al*., 2001). In addition, it will be important to assess the effect of mating rate. SR males have been shown to have smaller accessory glands, which act to reduce their mating frequency (Wilkinson, Swallow, Christensen, & Madden, 2003; Rogers, Denniff, Chapman, Fowler, & Pomiankowski, 2008; Meade, Finnegan, Kad, Fowler, & Pomiankowski, 2020). These further studies will provide a more comprehensive assessment of sperm competition as a factor contributing to the fertility of drive males and its consequences for the frequency of SR in wild populations.

In summary, we find no meiotic drive fertility cost under conditions of sperm competition, even though drive destroys half of carrier-male sperm. The absence of a fertility cost contributes to the relatively high frequency (∼20%) of meiotic drive in wild populations in this species (Wilkinson, Swallow, Christensen, & Madden, 2003; Cotton, Földvári, Cotton, & Pomiankowski, 2014; Paczolt, Reinhardt, & Wilkinson, 2017). This is unlike other species where drive males do poorly under multiple mating and the spread of drive is reliant on a high frequency of monandrous matings (Price, Hodgson, Lewis, Hurst, & Wedell, 2008; Courret, Chang, Wei, Montchamp-Moreau, & Larracuente, 2019; Dyer & Hall, 2019). The high fertility of drive males under sperm competition in *T. dalmanni* may reflect the way sperm is allocated in this polyandrous species, as multiple mating is a common feature amongst stalk-eyed fly species and precedes the evolution of meiotic drive in this clade (Wilkinson, Swallow, Christensen, & Madden, 2003). Alternatively, the absence of a fertility cost may be an evolved response to the loss of sperm caused by meiotic drive, which is supported by the finding male carriers of meiotic drive in this species have considerably enlarged testes (Meade, Finnegan, Kad, Fowler, & Pomiankowski, 2020).

## Acknowledgements

SB is supported by a Studentship from the Biotechnology and Biological Sciences Research Council (BB/M009513/1), AP is supported by funding from the Engineering and Physical Sciences Research Council (EP/F500351/1, EP/I017909/1), Natural Environment Research Council (NE/R010579/1) and Biotechnology and Biological Sciences Research Council (BB/V003542/1). We would like to thank Rebecca Finley, Wendy Hart and Matteo Mondani for maintaining the fly stocks and help with genotyping, and Kevin Fowler for advice on experimental design.

## Author Contributions

SB, LM and AP conceived the study. SB and LM carried out experiments and SB collected the data. SB, LM and AP analysed the data. SB and AP wrote the paper. All authors read, reviewed, and agreed on the submitted version.

## Data Archiving

The data that support the findings of this study are openly available in the Dryad at https://doi.org/10.5061/dryad.mkkwh713g.

