## Supplementary Material for "Meiotic drive does not impede success in sperm competition in the stalk-eyed fly, *Teleopsis dalmanni*"

Sade Bates, 0000-0002-9736-1077

Lara Meade, 0000-0002-5724-7413

Andrew Pomiankowski, 0000-0002-5171-8755

All statistical models and effect sizes

Supplementary Figures 1 and 2

Supplementary methods

### Supplementary information: Model Tables and Effect sizes

#### 1 Male fertility

##### 1.1 Variation in number of offspring sired with mating position

```
glm(formula = cbind(SR_offspring, ST_offspring) ~ as.factor(mating_position),
    family = quasibinomial, data = by_brood2)
```

Table 1: Analysis of variance table

| | Df | Deviance | Resid. Df | Resid. Dev | F | Pr(>F) |
|---|---|---|---|---|---|---|
| NULL | | | 44 | 503.358 | | |
| as.factor(mating_position) | 1 | 9.474 | 43 | 493.884 | 0.987 | 0.326 |

Table 2: Model Coefficients

| | Estimate | Std. Error | t value | Pr(>\|t\|) |
|---|---|---|---|---|
| (Intercept) | -0.137 | 0.267 | -0.513 | 0.611 |
| as.factor(mating_position)P2 | 0.373 | 0.376 | 0.991 | 0.327 |

The proportion of offspring sired by the P2 male was not different from 0.5 (mean P2 = 0.547 ± 0.0458) and there was no effect of mating position on the number of offspring sired per male.

##### 1.2 Variation in number of offspring sired with male genotype

```
glm(formula = cbind(P2_offspring, P1_offspring) ~ as.factor(male_genotype),
    family = quasibinomial, data = by_brood2)
```

Table 3: Analysis of variance table

| | Df | Deviance | Resid. Df | Resid. Dev | F | Pr(>F) |
|---|---|---|---|---|---|---|
| NULL | | | 44 | 494.551 | | |
| as.factor(male_genotype) | 1 | 0.666 | 43 | 493.884 | 0.069 | 0.793 |

Table 4: Model coefficients

| | Estimate | Std. Error | t value | Pr(>\|t\|) |
|---|---|---|---|---|
| (Intercept) | 0.236 | 0.265 | 0.890 | 0.378 |
| as.factor(male_genotype)ST | -0.099 | 0.376 | -0.263 | 0.793 |

The proportion of offspring sired by the SR male was not different from 0.5 (SR = 0.543 ± 0.046) and there was no effect of male genotype on the number of offspring sired per male.

#### 1.3 Variation in number of offspring sired with male genotype in the P1 role or P2 role

```
glm(formula = cbind(P1_offspring, P2_offspring) ~ as.factor(male_genotype),
    family = quasibinomial, data = P1_males_only)
```

Table 5: Analysis of variance table

| | Df | Deviance | Resid. Df | Resid. Dev | F | Pr(>F) |
|---|---|---|---|---|---|---|
| NULL | | | 45 | 512.502 | | |
| as.factor(male_genotype) | 1 | 1.471 | 44 | 511.031 | 0.152 | 0.699 |

Table 6: Model Coefficients

| | Estimate | Std. Error | t value | Pr(>\|t\|) |
|---|---|---|---|---|
| (Intercept) | -0.137 | 0.268 | -0.511 | 0.612 |
| as.factor(male_genotype)ST | -0.147 | 0.376 | -0.390 | 0.699 |

```
glm(formula = cbind(P1_offspring, P2_offspring) ~ as.factor(male_genotype),
    family = quasibinomial, data = P2_males_only)
```

Table 7: Analysis of variance table

| | Df | Deviance | Resid. Df | Resid. Dev | F | Pr(>F) |
|---|---|---|---|---|---|---|
| NULL | | | 45 | 500.679 | | |
| as.factor(male_genotype) | 1 | 0.157 | 44 | 500.522 | 0.016 | 0.899 |

Table 8: Model Coefficients

| | Estimate | Std. Error | t value | Pr(>\|t\|) |
|---|---|---|---|---|
| (Intercept) | -0.236 | 0.265 | -0.893 | 0.377 |
| as.factor(male_genotype)ST | 0.047 | 0.370 | 0.128 | 0.899 |

Examining the data from P1 males only, there was no effect of genotype on number of offspring sired. The same was true when examining data from P2 males only.

#### 1.4 Number of larvae collected and batch number

The number of larvae collected is a measure of female fecundity. A random sample of offspring from each female were genotyped. When the number of larvae collected and the batch were aby_male_ided as co-variates, neither mating order nor male genotype had an affect on the number of offspring sired by each male.

##### 1.4.1 Variation in P2 number of offspring sired with larvae collected and batch number

```
glm(formula = cbind(SR_offspring, ST_offspring) ~ batch + larvae_collected +
    mating_position, family = quasibinomial, data = by_brood2)
```

Table 9: Analysis of variance table (Type II tests)

| | Df | Deviance | Resid. Df | Resid. Dev | F | Pr(>F) |
|---|---|---|---|---|---|---|
| NULL | | | 44 | 503.358 | | |
| batch | 1 | 16.760 | 43 | 486.598 | 1.786 | 0.189 |
| larvae_collected | 1 | 21.192 | 42 | 465.405 | 2.258 | 0.141 |
| mating_position | 1 | 18.952 | 41 | 446.453 | 2.019 | 0.163 |

Table 10: Model Coefficients

| | Estimate | Std. Error | t value | Pr(>\|t\|) |
|---|---|---|---|---|
| (Intercept) | 0.259 | 1.014 | 0.256 | 0.800 |
| batch | 0.275 | 0.419 | 0.657 | 0.515 |
| larvae_collected | -0.016 | 0.009 | -1.728 | 0.092 |
| mating_positionP2 | 0.552 | 0.391 | 1.410 | 0.166 |

##### 1.4.2 Variation in SR number of offspring sired with larvae collected and batch number

```
glm(formula = cbind(P2_offspring, P1_offspring) ~ batch + larvae_collected +
    male_genotype, family = quasibinomial, data = by_brood2)
```

Table 11: Analysis of variance table (Type II tests)

| | Df | Deviance | Resid. Df | Resid. Dev | F | Pr(>F) |
|---|---|---|---|---|---|---|
| NULL | | | 44 | 494.551 | | |
| batch | 1 | 18.004 | 43 | 476.547 | 1.840 | 0.182 |
| larvae_collected | 1 | 0.201 | 42 | 476.346 | 0.021 | 0.887 |
| male_genotype | 1 | 1.003 | 41 | 475.343 | 0.103 | 0.750 |

Table 12: Model Coefficients

| | Estimate | Std. Error | t value | Pr(>\|t\|) |
|---|---|---|---|---|
| (Intercept) | -0.683 | 1.071 | -0.638 | 0.527 |
| batch | 0.552 | 0.425 | 1.298 | 0.202 |

| | Estimate | Std. Error | t value | Pr(>\|t\|) |
|---|---|---|---|---|
| larvae_collected | 0.001 | 0.009 | 0.078 | 0.939 |
| male_genotypeST | -0.125 | 0.392 | -0.320 | 0.751 |

#### 1.5 Male fertility and single parent broods

Table 13: Single parent broods

| mating position | proportion P2 offspring | male genotype | proportion SR offspring |
|---|---|---|---|
| P1 | 0.028 | ST | 0.972 |
| P1 | 0.000 | ST | 1.000 |
| P1 | 0.956 | ST | 0.044 |
| P2 | 0.955 | SR | 0.955 |
| P2 | 0.958 | SR | 0.958 |
| P1 | 0.967 | ST | 0.033 |
| P1 | 0.000 | ST | 1.000 |
| P2 | 0.960 | SR | 0.960 |
| P2 | 0.969 | SR | 0.969 |
| P2 | 0.034 | SR | 0.034 |
| P2 | 1.000 | SR | 1.000 |

##### 1.5.1 Variation in number of offspring sired with mating position with only single parent broods

```
glm(formula = cbind(SR_offspring, ST_offspring) ~ as.factor(mating_position),
    family = quasibinomial, data = subset(by_brood2, proportion_SR <=
        0.05 | proportion_SR >= 0.95))
```

Table 14: Analysis of variance table

| | Df | Deviance | Resid. Df | Resid. Dev | F | Pr(>F) |
|---|---|---|---|---|---|---|
| NULL | | | 10 | 300.235 | | |
| as.factor(mating_position) | 1 | 26.22 | 9 | 274.014 | 0.968 | 0.351 |

Table 15: Model Coefficients

| | Estimate | Std. Error | t value | Pr(>\|t\|) |
|---|---|---|---|---|
| (Intercept) | -0.042 | 0.871 | -0.048 | 0.963 |
| as.factor(mating_position)P2 | 1.295 | 1.359 | 0.953 | 0.366 |

When only extreme P2 offspring proportions of $\leq 0.05$ and $\geq 0.95$ are considered, the proportion of offspring sired by the P2 male was not different from 0.5 (proportion P2 = $0.621 \pm 0.145$), and mating order had no effect on the number of offspring sired.

##### 1.5.2 Variation in number of offspring sired with male genotype with only single parent broods

```
glm(formula = cbind(P2_offspring, P1_offspring) ~ as.factor(male_genotype),
    family = quasibinomial, data = subset(by_brood2, proportion_P2 <=
        0.05 | proportion_P2 >= 0.95))
```

Table 16: Analysis of variance table

| | Df | Deviance | Resid. Df | Resid. Dev | F | Pr(>F) |
|---|---|---|---|---|---|---|
| NULL | | | 10 | 296.799 | | |
| as.factor(male_genotype) | 1 | 22.784 | 9 | 274.014 | 0.841 | 0.383 |

Table 17: Model Coefficients

| | Estimate | Std. Error | t value | Pr(>\|t\|) |
|---|---|---|---|---|
| (Intercept) | 1.253 | 1.043 | 1.201 | 0.260 |
| as.factor(male_genotype)ST | -1.211 | 1.359 | -0.891 | 0.396 |

When only extreme SR offspring proportions of $\leq 0.05$ and $\geq 0.95$ are considered, the proportion of offspring sired by the SR male did not differ from 0.5 (proportion SR = $0.721 \pm 0.132$), and male genotype had no effect on number of offspring sired.

##### 1.5.3 Variation in number of offspring sired with mating position when single parent broods are excluded

```
glm(formula = cbind(SR_offspring, ST_offspring) ~ as.factor(mating_position),
    family = quasibinomial, data = subset(by_brood2, proportion_SR >
        0.05 & proportion_SR < 0.95))
```

Table 18: Analysis of variance table

| | Df | Deviance | Resid. Df | Resid. Dev | F | Pr(>F) |
|---|---|---|---|---|---|---|
| NULL | | | 33 | 179.949 | | |
| as.factor(mating_position) | 1 | 0.481 | 32 | 179.468 | 0.094 | 0.761 |

Table 19: Model Coefficients

| | Estimate | Std. Error | t value | Pr(>\|t\|) |
|---|---|---|---|---|
| (Intercept) | -0.171 | 0.227 | -0.753 | 0.457 |
| as.factor(mating_position)P2 | 0.098 | 0.319 | 0.307 | 0.761 |

When extreme P2 offspring proportions of $\leq 0.05$ and $\geq 0.95$ are excluded, the proportion of offspring sired by the P2 male was not different from 0.5 (proportion P2 = $0.523 \pm 0.04$) and mating order had no effect on number of offspring sired.

##### 1.5.4 Variation in male number of offspring sired with male genotype when single parent broods are excluded

```
glm(formula = cbind(P2_offspring, P1_offspring) ~ as.factor(male_genotype),
    family = quasibinomial, data = subset(by_brood2, proportion_P2 >=
        0.05 & proportion_P2 <= 0.95))
```

Table 20: Analysis of variance table

| | Df | Deviance | Resid. Df | Resid. Dev | F | Pr(>F) |
|---|---|---|---|---|---|---|
| NULL | | | 33 | 182.479 | | |
| as.factor(male_genotype) | 1 | 3.011 | 32 | 179.468 | 0.589 | 0.448 |

Table 21: Model Coefficients

| | Estimate | Std. Error | t value | Pr(>\|t\|) |
|---|---|---|---|---|
| (Intercept) | -0.073 | 0.224 | -0.328 | 0.745 |
| as.factor(male_genotype)ST | 0.245 | 0.319 | 0.767 | 0.449 |

When extreme SR offspring proportions of $\leq 0.05$ and $\geq 0.95$ are excluded, the proportion of offspring sired by the SR male was not different from 0.5 (proportion SR = $0.485 \pm 0.04$) and male genotype had no effect on number of offspring sired.

### 2 Male trait size and mating duration

#### 2.1 Variation in male traits

```
lm(formula = eyespan ~ thorax, data = by_male_id2)
```

Table 22: Analysis of variance table

| | Df | Sum Sq | Mean Sq | F value | Pr(>F) |
|---|---|---|---|---|---|
| thorax | 1 | 40.813 | 40.813 | 150.965 | 0 |
| Residuals | 88 | 23.791 | 0.270 | | |

Table 23: Model Coefficients

| | Estimate | Std. Error | t value | Pr(>\|t\|) |
|---|---|---|---|---|
| (Intercept) | -0.95 | 0.697 | -1.363 | 0.176 |
| thorax | 3.81 | 0.310 | 12.287 | 0.000 |

```
lm(formula = thorax ~ male_genotype, data = by_male_id2)
```

Table 24: Analysis of variance table

| | Df | Sum Sq | Mean Sq | F value | Pr(>F) |
|---|---|---|---|---|---|
| male_genotype | 1 | 0.169 | 0.169 | 5.644 | 0.02 |
| Residuals | 88 | 2.641 | 0.030 | | |

Table 25: Model Coefficients

| | Estimate | Std. Error | t value | Pr(>\|t\|) |
|---|---|---|---|---|
| (Intercept) | 2.198 | 0.026 | 84.142 | 0.00 |
| male_genotypeST | 0.087 | 0.037 | 2.376 | 0.02 |

```
lm(formula = eyespan ~ male_genotype, data = by_male_id2)
```

Table 26: Analysis of variance table

| | Df | Sum Sq | Mean Sq | F value | Pr(>F) |
|---|---|---|---|---|---|
| male_genotype | 1 | 5.915 | 5.915 | 8.868 | 0.004 |
| Residuals | 88 | 58.690 | 0.667 | | |

Table 27: Model Coefficients

| | Estimate | Std. Error | t value | Pr(>\|t\|) |
|---|---|---|---|---|
| (Intercept) | 7.331 | 0.123 | 59.544 | 0.000 |
| male_genotypeST | 0.513 | 0.172 | 2.978 | 0.004 |

```
lm(formula = eyespan ~ thorax + male_genotype, data = by_male_id2)
```

Table 28: Analysis of variance table (Type II tests)

| | Sum Sq | Df | F value | Pr(>F) |
|---|---|---|---|---|
| thorax | 35.692 | 1 | 135.027 | 0.000 |
| male_genotype | 0.794 | 1 | 3.002 | 0.087 |
| Residuals | 22.997 | 87 | | |

Table 29: Model Coefficients

| | Estimate | Std. Error | t value | Pr(>\|t\|) |
|---|---|---|---|---|
| (Intercept) | -0.748 | 0.700 | -1.069 | 0.288 |
| thorax | 3.676 | 0.316 | 11.620 | 0.000 |
| male_genotypeST | 0.194 | 0.112 | 1.733 | 0.087 |

```
lm(formula = eyespan ~ thorax:male_genotype, data = by_male_id2)
```

Table 30: Analysis of variance table

| | Sum Sq | Df | F value | Pr(>F) |
|---|---|---|---|---|
| thorax:male_genotype | 41.568 | 2 | 78.495 | 0 |
| Residuals | 23.036 | 87 | | |

Table 31: Model Coefficients

| | Estimate | Std. Error | t value | Pr(>\|t\|) |
|---|---|---|---|---|
| (Intercept) | -0.654 | 0.712 | -0.919 | 0.361 |
| thorax:male_genotypeSR | 3.635 | 0.324 | 11.214 | 0.000 |
| thorax:male_genotypeST | 3.719 | 0.312 | 11.931 | 0.000 |

Thorax length varies between male genotypes. Eyespan has strong covariance with thorax. Relative eyespan — the variation in eyespan not predicted by thorax length — is not significantly different between genotypes. Therefore, relative eyespan is not added to binomial GLMs for paternity.

#### 2.2 Male fertility with thorax length

```
glm(formula = cbind(ST_offspring, SR_offspring) ~ thorax.P1 +
    thorax.P2 + mating_position, family = quasibinomial, data = by_brood2)
```

Table 32: Analysis of variance table (Type II tests)

| | Sum Sq | Df | F value | Pr(>F) |
|---|---|---|---|---|
| thorax.P1 | 9.708 | 1 | 0.960 | 0.333 |
| thorax.P2 | 0.011 | 1 | 0.001 | 0.974 |
| mating_position | 7.567 | 1 | 0.748 | 0.392 |
| Residuals | 394.324 | 39 | | |

Table 33: Model Coefficients

| | Estimate | Std. Error | t value | Pr(>\|t\|) |
|---|---|---|---|---|
| (Intercept) | 2.665 | 3.634 | 0.733 | 0.468 |
| thorax.P1 | -1.098 | 1.126 | -0.975 | 0.336 |
| thorax.P2 | -0.043 | 1.315 | -0.033 | 0.974 |
| mating_positionP2 | -0.376 | 0.435 | -0.864 | 0.393 |

```
glm(formula = cbind(P1_offspring, P2_offspring) ~ thorax.ST +
    thorax.SR + male_genotype, family = quasibinomial, data = by_brood2)
```

Table 34: Analysis of variance table (Type II tests)

| | Sum Sq | Df | F value | Pr(>F) |
|---|---|---|---|---|
| thorax.ST | 0.721 | 1 | 0.071 | 0.792 |
| thorax.SR | 6.575 | 1 | 0.644 | 0.427 |
| male_genotype | 1.720 | 1 | 0.168 | 0.684 |
| Residuals | 398.274 | 39 | | |

Table 35: Model Coefficients

| | Estimate | Std. Error | t value | Pr(>\|t\|) |
|---|---|---|---|---|
| (Intercept) | -1.829 | 3.599 | -0.508 | 0.614 |
| thorax.ST | -0.311 | 1.170 | -0.266 | 0.792 |
| thorax.SR | 1.009 | 1.262 | 0.800 | 0.429 |
| male_genotypeST | 0.167 | 0.408 | 0.410 | 0.684 |

Though thorax length is different between SR and ST males, it does not affect number of offspring sired by each male, nor does it affect number of offspring sired by P2 and SR males.

#### 2.3 Male fertility with mating duration

Mating duration is the observed time taken for a single copulation in seconds.

##### 2.3.1 Variation in mating duration with mating position

```
lm(formula = mating_duration_sec ~ mating_position, data = by_male_id2)
```

Table 36: Analysis of variance table

| | Df | Sum Sq | Mean Sq | F value | Pr(>F) |
|---|---|---|---|---|---|
| mating_position | 1 | 3606.261 | 3606.261 | 1.38 | 0.243 |
| Residuals | 90 | 235147.348 | 2612.748 | | |

Table 37: Model Coefficients

| | Estimate | Std. Error | t value | Pr(>\|t\|) |
|---|---|---|---|---|
| (Intercept) | 63.674 | 7.537 | 8.449 | 0.000 |
| mating_positionP2 | 12.522 | 10.658 | 1.175 | 0.243 |

Mating duration did not differ between P1 and P2 males (mean P1 mating duration ± se: 64sec ± 3sec, mean P2 mating duration ± se: 76sec ± 3sec; $F_{1,90} = 1.38$, $P = 0.243$).

##### 2.3.2 Variation in mating duration with male genotype

```
lm(formula = mating_duration_sec ~ male_genotype, data = by_male_id2)
```

Table 38: Analysis of variance table

| | Df | Sum Sq | Mean Sq | F value | Pr(>F) |
|---|---|---|---|---|---|
| male_genotype | 1 | 6853.553 | 6853.553 | 2.66 | 0.106 |
| Residuals | 90 | 231900.056 | 2576.667 | | |

Table 39: Model Coefficients

| | Estimate | Std. Error | t value | Pr(>\|t\|) |
|---|---|---|---|---|
| (Intercept) | 78.756 | 7.567 | 10.408 | 0.000 |
| male_genotypeST | -17.266 | 10.587 | -1.631 | 0.106 |

Mating duration did not differ between ST and SR males (mean ST mating duration ± se: 61sec ± 3sec, mean SR mating duration ± se: 79sec ± 10sec; $F_{1,90} = 2.66$, $P = 0.106$).

##### 2.3.3 The effect of mating duration on the number of offspring sired per mating position

```
glm(formula = cbind(P2_offspring, P1_offspring) ~ mating_duration_sec.P1 +
    mating_duration_sec.P2, family = quasibinomial, data = by_brood2)
```

Table 40: Analysis of variance table

| | Sum Sq | Df | F value | Pr(>F) |
|---|---|---|---|---|
| mating_duration_sec.P1 | 31.954 | 1 | 3.474 | 0.069 |
| mating_duration_sec.P2 | 0.444 | 1 | 0.048 | 0.827 |

| | Sum Sq | Df | F value | Pr(>F) |
|---|---|---|---|---|
| Residuals | 386.285 | 42 | | |

Table 41: Model Coefficients

| | Estimate | Std. Error | t value | Pr(>\|t\|) |
|---|---|---|---|---|
| (Intercept) | 1.060 | 0.623 | 1.702 | 0.096 |
| mating_duration_sec.P1 | -0.015 | 0.008 | -1.760 | 0.086 |
| mating_duration_sec.P2 | 0.001 | 0.003 | 0.218 | 0.828 |

###### 2.3.4 The effect of mating duration on the number of offspring sired per male genotype

```
glm(formula = cbind(ST_offspring, SR_offspring) ~ mating_duration_sec.ST +
    mating_duration_sec.SR, family = quasibinomial, data = by_brood2)
```

Table 42: Analysis of variance table

| | Sum Sq | Df | F value | Pr(>F) |
|---|---|---|---|---|
| mating_duration_sec.ST | 39.549 | 1 | 4.292 | 0.044 |
| mating_duration_sec.SR | 1.732 | 1 | 0.188 | 0.667 |
| Residuals | 387.029 | 42 | | |

Table 43: Model Coefficients

| | Estimate | Std. Error | t value | Pr(>\|t\|) |
|---|---|---|---|---|
| (Intercept) | -1.638 | 0.961 | -1.705 | 0.096 |
| mating_duration_sec.ST | 0.028 | 0.015 | 1.915 | 0.062 |
| mating_duration_sec.SR | -0.001 | 0.003 | -0.427 | 0.671 |

###### 2.3.5 The effect of mating position with mating duration on the number of offspring sired

```
glm(formula = cbind(ST_offspring, SR_offspring) ~ mating_duration_sec.ST +
    mating_duration_sec.SR + mating_position, family = quasibinomial,
    data = by_brood2)
```

Table 44: Analysis of variance table (Type II tests)

| | Sum Sq | Df | F value | Pr(>F) |
|---|---|---|---|---|
| mating_duration_sec.ST | 39.917 | 1 | 4.245 | 0.046 |
| mating_duration_sec.SR | 0.545 | 1 | 0.058 | 0.811 |
| mating_position | 7.111 | 1 | 0.756 | 0.390 |
| Residuals | 385.517 | 41 | | |

Table 45: Model Coefficients

| | Estimate | Std. Error | t value | Pr(>\|t\|) |
|---|---|---|---|---|
| (Intercept) | -1.491 | 0.965 | -1.545 | 0.130 |
| mating_duration_sec.ST | 0.028 | 0.014 | 1.918 | 0.062 |
| mating_duration_sec.SR | -0.001 | 0.003 | -0.239 | 0.813 |
| mating_positionP2 | -0.338 | 0.389 | -0.868 | 0.390 |

Mating position still did not affect number of offspring sired by each male when mating duration was included as a covariate ($F_{1,41} = 0.756$, $P = 0.39$).

##### 2.3.6 The effect of male genotype with mating duration on the number of offspring sired

```
glm(formula = cbind(P1_offspring, P2_offspring) ~ mating_duration_sec.P1 +
    mating_duration_sec.P2 + male_genotype, family = quasibinomial,
    data = by_brood2)
```

Table 46: Analysis of variance table (Type II tests)

| | Sum Sq | Df | F value | Pr(>F) |
|---|---|---|---|---|
| mating_duration_sec.P1 | 31.780 | 1 | 3.373 | 0.074 |
| mating_duration_sec.P2 | 0.441 | 1 | 0.047 | 0.830 |
| male_genotype | 0.006 | 1 | 0.001 | 0.980 |
| Residuals | 386.268 | 41 | | |

Table 47: Model Coefficients

| | Estimate | Std. Error | t value | Pr(>\|t\|) |
|---|---|---|---|---|
| (Intercept) | -1.055 | 0.666 | -1.585 | 0.121 |
| mating_duration_sec.P1 | 0.015 | 0.008 | 1.737 | 0.090 |
| mating_duration_sec.P2 | -0.001 | 0.003 | -0.215 | 0.831 |
| male_genotypeST | -0.010 | 0.393 | -0.025 | 0.980 |

Male genotype type still does not affect the number of offspring sired by each male when mating during is included as a covariate ($F_{1,41} = 0.001$, $P = 0.98$).

#### 3 Supplementary figures

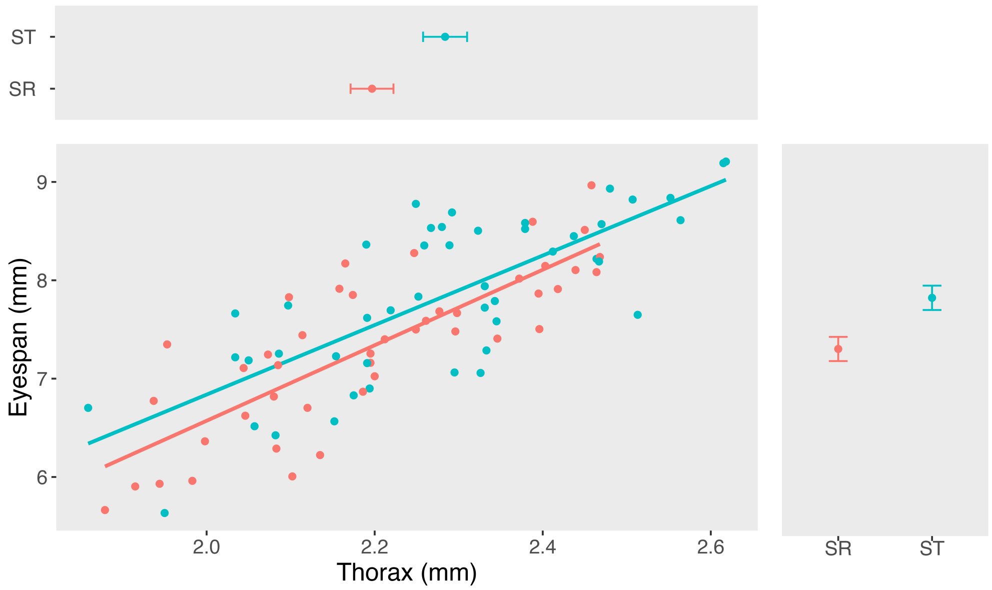


Figure 1: Variation in thorax length and eyespan with male genotype

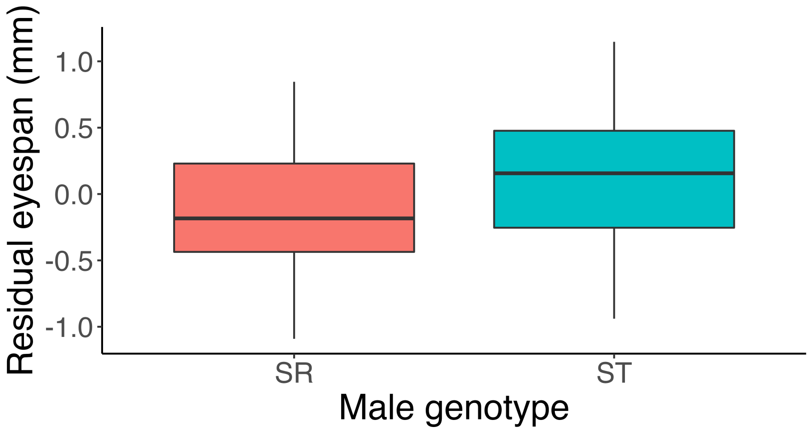


Figure 2: Variation in relative eyespan with genotype

**Supplementary Information: supplementary methods**

*The following protocol was used to extract and purify larval DNA (adapted from standard protocol in* Burke *et al.*, 1998)*:*

Larvae within each well were crushed using a micro-pestle prior to incubation for 16hrs at 55ºC, to extract DNA. The following day, 35µL 4M ammonium acetate was added to each sample to precipitate out proteins, and the plates chilled on ice for 5mins. The plates were then spun at 4450rpm, 4ºC for 60min. Next, the DNA was precipitated out by transferring 80µL of the supernatant from each sample to a new plate containing 80µL isopropanol per well. Centrifugation at 4450rpm, 4ºC for 60min pelleted out the DNA. The supernatant was discarded, and the DNA pellets were washed by adding 100µL 70% ethanol and spinning at 4450rpm for 30min. The ethanol was then removed, and the plates left to air dry for 1hr before adding 30µL T10 E0.1 buffer to each sample and incubating at 37ºC for 30mins to re-dissolve the DNA. Samples were stored at -20ºC prior to PCR analysis.

*The following PCR conditions were used for progeny genotyping:*

A 2720 Thermal Cycler (Applied Biosystems, Woolston, UK) was used to perform the reactions, which were carried out in 11µL volumes per sample, containing: 0.6µL forward and 0.6µL reverse primers (see Supp. Table 1 for sequences), both at 10µM, 0.12µL Phusion® High-Fidelity DNA Polymerase (New England Biolabs, Herts), 2.4µL Phusion® HF buffer (New England Biolabs, Herts), 6.4µL $ddH_20$ and either 1µL 10x diluted DNA, 5x diluted or pure DNA (depending on DNA concentration). The PCR programme was a 10min initial denaturation stage at 98ºC, followed by 45 cycles of 10sec denaturation at 98ºC, 30sec annealing time at 63ºC and 20sec extension at 72ºC. The reaction was completed by a 7min final extension step at 72ºC. The PCR products were analysed via gel electrophoresis on a 3% agarose/TBE gel run at 100V for ~1hr to separate them according to size and the results were visualised using a gel imaging system.

*Supplementary Table 1: comp16710 primer sequences*

| STRAND | Sequence |
|---|---|
| Forward | CGTGTCCGCATTTATACCAC |
| Reverse | GGTAGGCTTGTTCTAACGGC |
